# Integrating region- and network-level contributions to episodic recollection using multilevel structural equation modeling

**DOI:** 10.1101/2022.02.08.478511

**Authors:** Kyle A. Kurkela, Rose A. Cooper, Ehri Ryu, Maureen Ritchey

## Abstract

The brain is composed of networks of interacting brain regions that support higher order cognition. For instance, a posterior medial (PM) network appears to support recollection and other forms of episodic construction. Past research has focused largely on the roles of individual brain regions in recollection or on their mutual engagement as part of an integrated network. Yet the relationship between these region- and network-level contributions remains poorly understood. Here we applied multilevel structural equation modeling (SEM) to examine the functional organization of the PM network and its relationship to episodic memory outcomes. We evaluated two aspects of functional heterogeneity in the PM network: first, the organization of individual regions into subnetworks, and second, the presence of regionally-specific contributions while accounting for network-level effects. The results suggest that the PM network is composed of ventral and dorsal subnetworks, with the ventral subnetwork making a unique contribution to recollection, and that memory-related activity in individual regions is well accounted for by these network-level effects. These findings highlight the importance of considering both region and network levels of analysis when studying brain-behavior relationships.

---

The brain is composed of networks of interacting brain regions, and the functioning of these networks is crucial for higher-level cognition. One network of brain regions in particular, the posterior medial (PM) network, has consistently been linked to our ability to mentally construct events – recollecting past events, imaging future events, or constructing fictional scenarios (Buckner & DiNicola, 2019; Ritchey & Cooper, 2020). Exactly how this network supports episodic construction, however, remains unclear. Past research suggests that there are both regional and network level contributions of the PM network to episodic recollection. For instance, the hippocampus has long been known to be essential for episodic memory (e.g., Corkin, 2002; Riedel et al., 1999) and neuroimaging studies have shown that its activity is correlated with successful recollection (H. Kim, 2013; Spaniol et al., 2009). Looking beyond the hippocampus, however, it is apparent that many other brain regions are also involved in the recollection process (H. Kim, 2013; Rugg & Vilberg, 2013; Spaniol et al., 2009) including the angular gyrus (AG), precuneus (Prec), retrosplenial cortex (RSC), posterior cingulate cortex (PCC), parahippocampal cortex (PHC), and medial prefrontal cortex (MPFC). These regions are robustly structurally and functionally connected with the hippocampus, leading to proposals that they form an integrated network (Buckner & DiNicola, 2019; Ranganath & Ritchey, 2012; Rugg & Vilberg, 2013). Here we use multilevel structural equation modeling to examine heterogeneity in the function of the PM network during an episodic retrieval task. Specifically, we investigated the subnetwork architecture of the PM network, as well as the contributions of individual PM regions to predicting memory outcomes.

A great deal of research has focused on the roles of individual brain regions in supporting episodic construction, delineating specific roles for the hippocampus, angular gyrus, and other regions of the PM network (Ritchey & Cooper, 2020). The hippocampus, for example, is posited to support the binding together of contextual details in memory (Davachi, 2006; Diana et al., 2007; Eichenbaum et al., 2007) and is thought to perform a pattern completion function in which partial representations evoked by memory cues are “completed” by reinstating related information stored in memory (Horner et al., 2015; Marr, 1971; Norman & O’Reilly, 2003). The angular gyrus, on the other hand, is thought to support the representation of multimodal episodic details brought to mind during recollection (Humphreys et al., 2021; Ramanan et al., 2018; Rugg & King, 2018). Some fMRI studies have directly tested for cognitive and temporal dissociations amongst the different regions of the PM network, finding evidence for functional specialization in the context of both episodic memory (Richter et al., 2016; Vilberg & Rugg, 2012, 2014) and imagination (Thakral et al., 2020). For example, Richter and colleagues (2016) used fMRI to identify brain activity that tracked the success, precision, and subjective vividness of episodic recollection. The authors modeled these measures jointly and found that the hippocampus uniquely tracked whether or not retrieval was successful, the angular gyrus uniquely tracked the precision of remembered information, and the precuneus uniquely tracked subjective memory vividness. These findings suggest that individual regions of the PM network make distinct contributions to the recollection process.

There is other evidence pointing to the importance of mutual engagement of these regions as part of an integrated network. Functional MRI studies have consistently observed increased neural activity across the PM network that is related to successful, vivid episodic recollection (H. Kim, 2013; Rugg & Vilberg, 2013; Spaniol et al., 2009). Moreover, functional connectivity within the PM network scales with the quality of recollection, including its subjective vividness or the recovery of source details (Cooper & Ritchey, 2019; Geib, Stanley, Dennis, et al., 2017; Geib, Stanley, Wing, et al., 2017; King et al., 2015; Schedlbauer et al., 2014; Watrous et al., 2013). Interestingly, another line of research suggests that the PM network may not act as a single homogeneous network but is instead composed of at least two, highly related subnetworks (Andrews-Hanna et al., 2010; Barnett et al., 2021; Buckner & DiNicola, 2019; Cooper et al., 2021). For example, Cooper and colleagues (2021) examined the functional connectivity of the PM network regions while participants watched a short movie and, using data-driven community detection, showed that the PM network fractured into ventral and dorsal subnetworks that were differentially engaged during event perception. Barnett and colleagues (2021) used a similar data-driven approach to delineate multiple subnetworks in the PM network using resting-state data while also showing evidence that the regions of the different PM subnetworks represented similar information during a memory-guided decision making task. While these studies provide support for a subnetwork view of the PM network, it currently remains unclear whether these subnetworks are dissociable in their contributions to episodic recollection.

Prior studies have made it clear that activity in regions of the PM network are related to memory and to one another, but it remains unclear whether their contributions to memory are regionally specific or shared across the network. Structural equation modeling (SEM) is a well suited tool for delineating regional and network-level contributions to behavior (Bolt et al., 2018). SEM allows for the capturing of a common, distributed, network-level contribution by estimating latent variables that captures the covariance amongst regions. Structural models can then estimate the statistical dependency between these latent variables and some behavioral variable while also estimating the regional-specific effect of each of the regions, statistically controlling for their membership within larger networks. For instance, Bolt and colleagues (2018) used SEM to parse the unique contributions of the right dorsolateral prefrontal cortex to cognitive control from those of the larger frontoparietal control network, showing that the unit of behavioral significance for many common cognitive control tasks was not the right dorsolateral prefrontal cortex, but the shared contributions of the frontoparietal control network. In the context of episodic memory, we can use similar tools to estimate the specific regional contributions and common network contributions to episodic remembering within a single statistical model.

Taken together, the literature suggests that the regions of the PM network perform dissociable yet interrelated functions and, as a result, make separable contributions to the recollection process. It remains unclear, however, exactly how to combine the findings from experiments taking region-focused approaches and network-focused approaches — highlighting the need for an approach that can simultaneously take into account network-wide and region-specific contributions to episodic retrieval. The present study uses SEM to model heterogeneity of function of the PM network. Specifically, we seek to model two key aspects of functional heterogeneity within the network: that larger networks fracture into related subnetworks and that regions of the networks make extra-network contributions to cognition.

## Methods

### Experiment

#### Participants

Twenty eight participants from Cooper & Ritchey (2019) were included in the final set of analyses after excluding participants who did not complete the study or who had inadequate memory performance (see Cooper & Ritchey 2019). Participants were selected such that they were between the ages of 18 and 35 (*M* = 21.82, *SD* = 3.57, 16 females, 12 males) and had no history of neurological or psychiatric illness. Informed consent was obtained from all participants prior to the experiment and participants were reimbursed for their time. All procedures were approved by the Boston College Institutional Review Board.

## Materials

Memoranda consisted of 144 unique events that were constructed using a combination of 144 episode-unique object stimuli from Brady and colleagues (2013), 6 panoramic scenes from the SUN 360 database (Xiao et al., 2012), and 12 sounds from the International Affective Digitized Sounds (IADS) database (Bradley & Lang, 2007). See Cooper & Ritchey (2019) for further information.

### Procedure

Participants completed six interleaved encoding and retrieval phases while undergoing MRI scanning (see Cooper & Ritchey, 2019). To summarize, during scanned encoding phases participants were told that they would encounter 24 events consisting of a foreground object, a background scene, and an emotionally evocative sound. Participants were asked to remember each of the events in as much detail as possible, with explicit instructions to try and remember the color of the foreground object, the position of the object within the background scene, and the emotional valence of the sound. During scanned retrieval phases, participants were tested on their ability to reconstruct episode features from memory. At the beginning of each retrieval trial, participants were shown grayscale versions of the object stimuli from the previous encoding phase. During this time, participants were asked to bring to mind the cued episode in as much detail as possible. Immediately after this remember period, participants were asked to report the emotional valence of the episode’s sound using a confidence scale. The confidence scale asked participants to identify their response to the emotional valence question as either with high confidence or with low confidence. After reporting the sound’s valence, participants were asked to report the quality of their memory for the remaining two features in a counterbalanced order. Specifically, participants were presented with the object image in a random color on top of a random view from the correct background scene. Participants were instructed to reconstruct the color of the target image using an interactive 360-degree color wheel and to position the object within the background scene using a similar interactive 360-degree panoramic scale.

### FMRI data acquisition

MRI scanning was performed at the Harvard Center for Brain Science using a 3T Siemens Prisma MRI scanner with a 32-channel head coil. Structural MRI images were collected using a T-1 weighted multiecho MPRAGE protocol with a field of view = 256 mm, 1 mm isotropic voxels, 176 sagittal slices with an interleaved acquisition, TR = 2530 ms, TE = 1.69/3.55/5.41/7.27 ms, flip angle = 7 degrees, phase encoding from anterior-posterior, parallel imaging = GRAPPA, and an acceleration factor of 2. Functional images were acquired using a whole brain multiband echo-planar imaging sequence with a field of view of 208 mm, 2 mm isotropic voxels, 69 slices at T > C -25.0 with interleaved acquisition, TR = 1500 ms, TE = 28 ms, flip angle = 75 degrees, anterior-posterior phase encoding, parallel imaging with GRAPPA, and an acceleration factor of 2. A total of 6 scan runs were collected, each of which consisted of 466 TRs.

## Analyses

### Behavioral Data

Behavioral data from the dataset consisted of trialwise error values measured in degrees for the object color and scene position features and of binary data (i.e., 1: correct; 0: incorrect) for the sound valence feature (i.e., collapsed across confidence). For consistency across behavioral measures, we transformed the object color and scene position measures into binary measures representing whether a response was correct (1) or incorrect (0), similar to what we have done previously (Cooper & Ritchey 2019). This was done by taking any trial with an error smaller than the accuracy threshold (see below) and giving it a score of 1 and taking any trial with an error greater than the accuracy threshold and giving it a score of 0. Descriptive statistics of our behavioral variables are detailed in Table 1.

**Table 1:** Descriptive statistics for variables of interest. Neural measures (pHipp-pAG) are the mean t-value across all voxels within that ROI from the single-trial estimation step. Behavioral measures (SCENE, COLOR, SOUND) are binary, coded such that 1 = correct and 0 = incorrect. Correlations between neural measures are Pearson’s Correlation Coefficients. Correlations between the behavioral and neural variables are Point-Biserial Correlations Coefficients. All descriptive statistics, excluding the ICCs, were calculated ignoring the nested structure. pHipp = posterior hippocampus, Prec = precuneus, PCC = posterior cingulate cortex, MPFC = medial prefrontal cortex, PHC = parahippocampal cortex, RSC = retrosplenial cortex, aAG = anterior angular gyrus, pAG = posterior angular gyrus, SCENE = Scene Position Correct, COLOR = Object Color Correct, SOUND = Sound Valence Correct, sd = standard deviation, ICC = interclass correlation coefficient.

|  | mean | sd | min | max | ICC | Correlations |  |  |  |  |  |  |  |  |  |
| --- | --- | --- | --- | --- | --- | --- | --- | --- | --- | --- | --- | --- | --- | --- | --- |
|  |  |  |  |  |  | RSC | PHC | pAG | aAG | pHipp | Prec | PCC | MPFC | SCENE | COLOR |
| RSC | 0.049 | 0.170 | -0.567 | 0.808 | 0.138 |  |  |  |  |  |  |  |  |  |  |
| PHC | -0.028 | 0.181 | -0.763 | 1.048 | 0.155 | 0.324 |  |  |  |  |  |  |  |  |  |
| pAG | -0.007 | 0.315 | -1.292 | 1.232 | 0.138 | 0.296 | 0.330 |  |  |  |  |  |  |  |  |
| aAG | 0.188 | 0.352 | -1.346 | 1.648 | 0.184 | 0.296 | 0.119 | 0.295 |  |  |  |  |  |  |  |
| pHipp | 0.077 | 0.156 | -0.614 | 1.182 | 0.022 | 0.191 | 0.184 | 0.135 | 0.276 |  |  |  |  |  |  |
| Prec | 0.116 | 0.215 | -0.923 | 0.971 | 0.123 | 0.326 | 0.265 | 0.247 | 0.356 | 0.285 |  |  |  |  |  |
| PCC | 0.144 | 0.218 | -0.614 | 1.044 | 0.142 | 0.282 | 0.173 | 0.189 | 0.418 | 0.283 | 0.410 |  |  |  |  |
| MPFC | 0.135 | 0.279 | -1.176 | 1.308 | 0.036 | 0.162 | 0.101 | 0.066 | 0.342 | 0.285 | 0.282 | 0.402 |  |  |  |
| SCENE | 0.675 | 0.468 | 0.000 | 1.000 | 0.205 | 0.194 | 0.130 | 0.086 | 0.050 | 0.051 | 0.094 | 0.055 | -0.026 |  |  |
| COLOR | 0.724 | 0.447 | 0.000 | 1.000 | 0.135 | 0.070 | 0.046 | 0.024 | 0.046 | 0.032 | 0.081 | 0.061 | -0.033 | 0.251 |  |
| SOUND | 0.758 | 0.429 | 0.000 | 1.000 | 0.118 | 0.091 | 0.070 | 0.034 | 0.039 | 0.066 | 0.099 | 0.078 | 0.021 | 0.229 | 0.170 |

The accuracy threshold was determined by fitting two probability density functions to group aggregate data within a mixture modeling framework and is described in detail in Cooper and Ritchey (2019). In brief, Cooper and Ritchey (2019) estimated (for the color and scene features separately) the probability that each error resulted from the von Mises as opposed to the uniform distribution. Errors that had less than a 50% chance of fitting the von Mises distribution were labeled as incorrect. This analysis resulted in a threshold of +/- 57 degrees for the object color feature and +/- 30 degrees for the scene position feature and served as our threshold for labeling a response as “correct” or “incorrect”.

### MRI Preprocessing

All preprocessing of the MRI data was performed using FMRIPrep v1.0.3 (Esteban et al., 2019). Data were preprocessed using the same steps as in Cooper & Ritchey (2019). First, each T1w volume was corrected for intensity non uniformity and skull-stripped. Spatial normalization to the ICBM 152 Nonlinear Asymmetrical template version 2009c was performed through nonlinear registration, using brain-extracted versions of both the T1w volume and template. All analyses reported here use structural and functional data in MNI space. Brain tissue segmentation of cerebrospinal fluid (CSF), white-matter (WM), and gray-matter (GM) was performed on the brain-extracted T1w image. Functional data was slice time corrected, motion corrected, and corrected for field distortion. This was followed by co-registration to the corresponding T1w using boundary-based registration with 9 degrees of freedom. Physiological noise regressors were extracted using CompCor. A mask to exclude signals with cortical origin was obtained by eroding the brain mask, ensuring it only contained subcortical structures. Six aCompCor components were calculated within the intersection of the subcortical mask and the union of CSF and WM masks calculated in T1w space, after their projection to the native space of each functional run. Framewise displacement was also calculated for each functional run. No smoothing of the data was performed. For further details of the pipeline please refer to the online documentation: https://fmriprep.readthedocs.io/en/1.0.3/index.html.

### Trialwise Response Estimates

To estimate trialwise response estimates, we next used a multi-model approach proposed by Mumford and colleagues (2012) and implemented in SPM12 (https://www.fil.ion.ucl.ac.uk/spm/) using in house MATLAB (https://www.mathworks.com/products/matlab.html) scripts. In this approach, a separate general linear model is built to estimate the amplitude of the BOLD response for each trial by modeling the response for each trial using its own dedicated regressor and modeling all other trials as a separate regressor (including the following nuisance regressors: translation in the x, y, and z dimensions; rotation in pitch, roll, and yaw; the first principal component from aCompCorr, and framewise displacement). The GLMs for the current study were limited to TRs from the retrieval phase from Cooper and Ritchey (2019). Each retrieval trial was modeled by convolving SPM12’s hemodynamic response function with a stick function placed at the onset of each retrieval trial.The statistic used as our estimate of the BOLD response was the t-statistic for the regressor corresponding to each individual trial. The t-statistic provides a more sensitive measure than beta values when searching for information within the brain (Misaki et al., 2010) and downweights noisy voxels, allowing them to have a smaller influence on results Trialwise response estimates were extracted from our regions of interest (ROIs) and submitted to further analysis.

### ROIs

The ROIs for the present analysis are the same ones used in a previous study from our lab investigating interactions among PM regions (Cooper et al., 2021). The ROIs were created using a combination of cortical ROIs from the Schaefer Atlas (Schaefer et al., 2018) and a hippocampal ROI from a probabilistic parcellation (Ritchey et al., 2015). These anatomical ROIs were combined with a meta-analytic map generated using Neurosynth (Yarkoni et al., 2011) using the search term “episodic”. Functional peaks from this map within regions of the PM network were selected and ROIs were drawn around these peaks such that they were of equal size and each contained 100 contiguous voxels. The final set of ROIs included the posterior hippocampus (pHipp), the parahippocampal cortex (PHC), the retrosplenial cortex (RSC), the precuneus (Prec), posterior cingulate cortex (PCC), posterior angular gyrus (pAG), anterior angular gyrus (aAG), and the medial prefrontal cortex (MPFC).

### Multilevel Structural Equation Modeling

The present study took a multilevel structural equation modeling (SEM) approach to investigate functional heterogeneity in the PM network. Multilevel SEM allows for the estimation of latent constructs and for modeling the structural paths amongst those latent constructs in datasets that have a nested structure. This approach is optimal for the current dataset that contains observations of ROI activity across trials that are nested within subjects. In our data, trials are the level-1 units and subjects are the level-2 units. Because of the nested structure, the variables that vary at the trial level have two sources of variation: one due to the difference between trials within subjects and the other due to the difference between subjects.For the neural data, the former represents where the BOLD response estimate (i.e., the t value) is relative to that subject’s own average across all trials and the latter represents where each subject’s average t value compared to other subjects’ average t values. Our primary interest was in modeling within-subject, trial-to-trial variability. The between-subject variability in the neural data could represent meaningful differences in individual characteristics, but we did not have

*a-priori* hypotheses about these individual differences in the present sample. Therefore, in the multilevel model for the neural data (see below), the between-subject model is specified only so that this source of variability is accounted for and therefore the statistical validity of the within-subject model is not compromised. For the behavioral data, the two sources of variation represent differences in overall memory quality on a trial-to-trial basis and differences in overall accuracy across trials on a subject-to-subject basis.

All modeling was performed using *Mplus* software version 8.2 (Muthén & Muthén,1997-2017). Models were determined to have acceptable levels of model fit if they displayed the following fit indices: root mean squared error of approximation (RMSEA) < .08, comparative fit index (CFI) > .95, and a standardized root mean squared residual (SRMR^1^) < .06 (Hu & Bentler,1999). For SRMR, a separate index was calculated for the within cluster (i.e., within subject) and between cluster (i.e., between subject) levels. The models were estimated using the MLR (maximum likelihood with robust standard errors) and the WLSMV (weighted least squares means and variances adjusted) estimators in *Mplus*.

### Preliminary Analyses

Prior to performing our multilevel SEM analysis, we verified the necessity of a multilevel analysis by calculating intraclass correlations (ICCs) for each of our variables of interest and by fitting a “null” model designed to test if there is any structure in the between-subjects covariance matrix (see Jak et al., 2013). The ICC is a statistic that reflects the proportion of variance of a variable that can be attributed to individual differences amongst our subjects. Datasets that contain variables that have ICCs close to 0 will not see an additional benefit from multilevel modeling. In the null model, a saturated (i.e., a model that estimates parameters for all possible variances and covariances amongst variables) was specified for the within-subject covariance structure, and a *null* model in which all of the variances and covariances are constrained to be zero was specified for the between-subject covariance structure. If this model fails to satisfactorily fit the data, it suggests that the between-subject variances need to be allowed in the model and therefore calls for a two-level model. If the ICCs are greater than 0.1 or the null model fits the data poorly, we will conclude that multilevel modeling is required for our dataset.

### Measurement Models

After verifying the necessity of multilevel modeling, we examined the measurement structure underlying our eight ROIs. We tested two measurement models: a single-factor model for the integrated PM network hypothesis and a two factor model for the two subnetwork hypothesis. We tested two measurement models: a single-factor model for the integrated PM network hypothesis and a two factor model for the two subnetworks hypothesis. In the single-factor model, the eight PM network regions loaded onto a single latent factor at the within-subject level. We did not impose any restriction at the between-subject level because we did not have a-priori hypotheses about the nature of the between-subject variability in neural data. In the two factor model, RSC, PHC, and pAG loaded on one factor representing the ventral Posterior Medial Network (vPMN), and MPFC, pHipp, Prec, PCC, and aAG loaded on the other factor representing the dorsal Posterior Medial Network (dPMN) (see Figure 1). Again, we did not impose any restriction at the between-subject level. For the measurement models for neural data, the MLR estimator was used.

**Figure 1:**
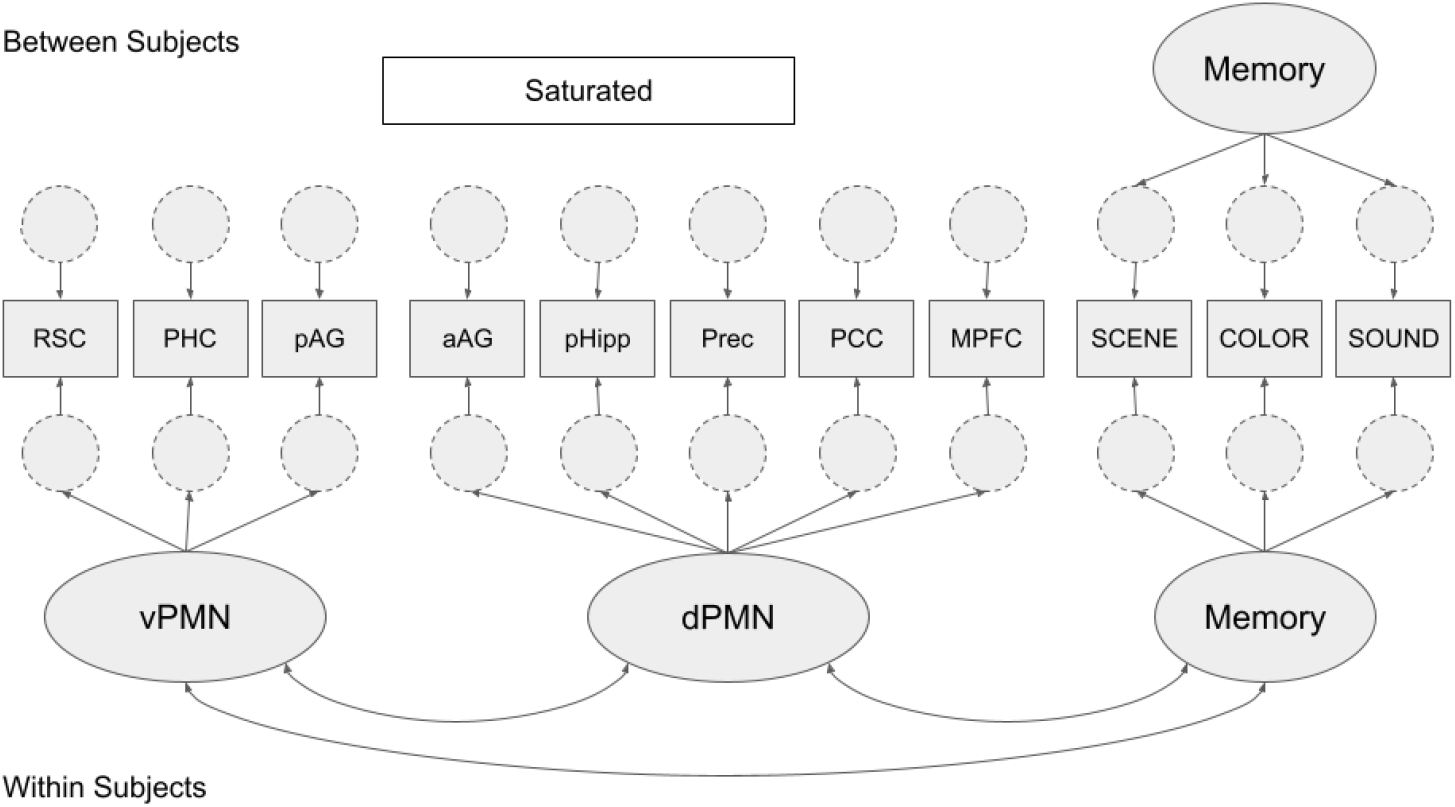
Measurement Model. A graphical representation of our measurement model following the graphing conventions of E. Kim and colleagues (2016). Our measurement model contained two latent variables for the neural data at the within-subjects level, a single latent variable for the behavioral data at the within-subjects level, and a single latent variable representing the behavioral data at the between-subjects level. The factor loadings for the Memory latent variables were set equal across the levels. At the between subject level, the eight neural variables were allowed to covary with one another and with the Memory factor. See Table 2 for standardized parameters of the within-subjects part of the model. vPMN = ventral posterior medial network, dPMN = dorsal posterior medial network, Memory = overall memory quality, pHipp = posterior hippocampus, Prec = precuneus, PCC = posterior cingulate cortex, MPFC = medial prefrontal cortex, PHC = parahippocampal cortex, RSC = retrosplenial cortex, aAG = anterior angular gyrus, pAG = posterior angular gyrus, Scene = scene position feature correct, Color = object color feature correct, Sound = sound valence feature correct.

For the behavioral data, we fit a two-level categorical factor model with a single factor specified at both levels, such that all three behavioral variables representing memory for different aspects of our events (i.e., Scene Position, Object Color, Sound Valence) loaded onto a single latent variable. Because the behavioral variables were binary, the WLSMV estimator was used.The latent factor at the within-subjects level represents the overall memory quality on a trial-to-trial basis. The latent factor at the between-subjects level represents participants’ overall memory ability. After fitting this initial model, we tested for metric invariance (i.e., equal factor loadings across levels) in this two-level factor model. If metric invariance does hold, we can interpret the between-subjects latent factor as the random intercept of the within-subject latent factor and calculate the ICC for the memory quality factor.

### Structural Models

After establishing good-fitting measurement models, the neural and behavioral models were stitched together to form a single model (see Figure 1). We subsequently fit a series of models to the data to quantify the contribution of the PM network (or PM subnetworks) to memory quality and whether any of the regions made a region-specific contribution to memory quality over and above their network (or subnetwork) contribution. For these models, the WLSMV estimator was used. A baseline structural model (*Model 0*) contained a structural path from the PM Network (or PM subnetworks) latent variable(s) to the Memory Quality latent variable at the within-subjects level. After fitting the baseline model, a series of models were fit to test for region-specific contributions (i.e., one at a time). Each of these models included an additional structural path from the region to Memory Quality. This direct path reflects the predictive effect of the region after accounting for its participation in the network (or subnetworks). Wald’s test statistic was used to determine the statistical significance of the additional path.

## Results

### Preliminary Analyses

The ICCs were larger than 0.1 for the neural measures, except for pHipp whose ICC was .022 and MPFC whose ICC was .036. The null model for the neural data did not fit the data well (*χ*^2^ = 2951.578, df = 36, *p* < .001, RMSEA = 0.144, CFI = 0.000, SRMRwithin = 0.047, SRMRbetween= 0.352, AIC = -10496.752, BIC = -10221.064). Thus, we concluded that multilevel modeling was appropriate for our neural data. For the behavioral measures, the ICCs ranged from .118 to .205, indicating that about 10 to 20% of the variance in the memory measures are due to between-subjects differences. The null model resulted in adequate fit to the data (*χ*^2^ = 20.882, df = 6, *p* < .01, RMSEA = 0.025, CFI = 0.958, SRMRwithin = 0.000, SRMRbetween= 0.634).The fit statistics for the behavioral null model were obtained with the WLSMV estimator. Applying the same criteria for the WLSMV fit statistics have been shown to be less sensitive to discover model-data misfit (Xia & Yang, 2019). Considering the large ICC values and the limitation in the performance of the WLSMV fit statistics, we concluded that adopting a multilevel model was appropriate for the behavioral data.

### Measurement Models

The one factor model for the neural data resulted in the following fit statistics (*χ*^2^ = 290.442, df = 20, *p* < .001, RMSEA = 0.059, CFI = 0.905, SRMRwithin = 0.063, SRMRbetween=0.005, *AIC* = -12584.469, *BIC* = -12208.530). The two factor model fit the data well and better than the one factor model (*χ*^2^ = 90.096, df = 19, *p* < .001, RMSEA = 0.031, CFI = 0.975, SRMRwithin = 0.035, SRMRbetween= 0.003, *AIC* = -13145.343, *BIC* = -12763.138). To further compare model fits, we examined the estimated correlation between the two latent factors in the two factor model and compared the estimated communality values of the two models. We reasoned that additional evidence in favor of the two factor model would be seen if the correlation between the latent factors was estimated to be low-moderate *and* if the estimated communality values were all equivalent or higher for the two factor model relative to the one factor model. We note that the correlation between the vPMN and dPMN latent variables was high, but not perfect (*r* = 0.630 or ∼39.7% of variance shared) and the communality values in the two factor model were all equivalent or higher compared with the one factor model (see Table 2). Taken together, these results suggest that a two factor model was a better model for our neural data.

**Table 2:** Communality Values. Communality value estimates from the One Factor and Two Factor measurement models. All of the estimated communality values are equivalent or higher in the Two Factor model compared with the One Factor model. param = parameter, est = estimate se = standard error, pval = p value. pHipp = posterior hippocampus, Prec = precuneus, PCC = posterior cingulate cortex, MPFC = medial prefrontal cortex, PHC = parahippocampal cortex, RSC = retrosplenial cortex, aAG = anterior angular gyrus, pAG = posterior angular gyrus.

| param | One Factor |  |  | Two Factor |  |  |
| --- | --- | --- | --- | --- | --- | --- |
|  | est | se | pval | est | se | pval |
| pHipp | 0.188 | 0.024 | < 0.001 | 0.193 | 0.025 | < 0.001 |
| Prec | 0.397 | 0.039 | < 0.001 | 0.395 | 0.039 | < 0.001 |
| PCC | 0.437 | 0.029 | < 0.001 | 0.479 | 0.024 | < 0.001 |
| MPFC | 0.251 | 0.030 | < 0.001 | 0.286 | 0.032 | < 0.001 |
| PHC | 0.177 | 0.027 | < 0.001 | 0.344 | 0.030 | < 0.001 |
| RSC | 0.231 | 0.036 | < 0.001 | 0.391 | 0.035 | < 0.001 |
| aAG | 0.472 | 0.030 | < 0.001 | 0.485 | 0.032 | < 0.001 |
| pAG | 0.215 | 0.038 | < 0.001 | 0.363 | 0.036 | < 0.001 |

The two-level single-factor model for the behavioral data was just identified (i.e., df = 0) and thus fit the data perfectly since the number of covariances (12 - 6 at the between-subjects level and 6 at the within-subject level) exactly equals the number of free parameters (12). The model with equality constraints on the factor loadings for the metric invariance across levels fit the data well (*χ*^2^ = 0.163, df = 2, *p* < .9218, RMSEA = 0.000, CFI = 1.00, SRMRwithin = 0.001, SRMRbetween= 0.022). With metric invariance, the latent variables at each level can be interpreted as the within-subject and between-subject components, respectively, of the same construct “overall memory quality”. Metric invariance additionally allowed us to calculate the proportion of variance in the overall memory quality factor that is attributable to individual differences and trial-to-trial differences. Overall, the memory quality factor had an ICC of .329, meaning that individual differences explained approximately 32.9% of the variability in memory quality. This is advantageous for our purposes, since our primary interest was explaining trial-to-trial variability in memory quality.

After finding good fitting neural and behavioral measurement models, we proceeded to fit a joint measurement model by stitching the two factor neural model and the metric-invariance behavioral model together (see Figure 1). At the between subjects level, the regions were allowed to covary with one another and also allowed to covary with the between-subject memory quality factor. This joint measurement model fit the data adequately (*χ*^2^ = 534.782, df = 59, *p* <.001, RMSEA = 0.046, CFI = 0.974, SRMRwithin = 0.034, SRMRbetween= 0.043). The key parameter estimates for the within-subject part of this model are reported in Table 3.

**Table 3:** Measurement Model Standardized Parameter Estimates. Select standardized parameter estimates in the within-subject level model. This table was created using the R package *MplusAutomation* (Hallquist & Wiley, 2018). Parameter headers (paramHeader) follow standard Mplus syntax, where the BY keyword indicates a loading parameter (lambda λ) and the WITH keyword indicates a covariance parameter (theta θ). param = parameter, est = estimate, se = standard error, pval = p value. See Figure 1 caption for abbreviations.

| paramHeader | param | est | se | pval |
| --- | --- | --- | --- | --- |
| vPMN.BY | RSC | 0.627 | 0.006 | < 0.001 |
|  | PHC | 0.567 | 0.008 | < 0.001 |
|  | pAG | 0.608 | 0.007 | < 0.001 |
| dPMN.BY | MPFC | 0.516 | 0.006 | < 0.001 |
|  | pHipp | 0.437 | 0.008 | < 0.001 |
|  | Prec | 0.647 | 0.007 | < 0.001 |
|  | PCC | 0.685 | 0.004 | < 0.001 |
|  | aAG | 0.711 | 0.005 | < 0.001 |
| Memory.BY | SCENE | 0.707 | 0.057 | < 0.001 |
|  | COLOR | 0.465 | 0.044 | < 0.001 |
|  | SOUND | 0.460 | 0.030 | < 0.001 |
| dPMN.WITH | vPMN | 0.634 | 0.008 | < 0.001 |
| Memory.WITH | vPMN | 0.200 | 0.025 | < 0.001 |
|  | dPMN | 0.102 | 0.026 | < 0.001 |

### Structural Models

We estimated a series of structural models by specifying structural relationships among the latent factors in the joint measurement model. The fit statistics are presented in Table 3. In the first model (*Model 0: Baseline*, see Figure 2), each of the two subnetworks was allowed to have a structural path to memory quality. In this baseline model, the vPMN uniquely (i.e., when statistically controlling for the dPMN) predicted the overall quality with which events were remembered (*β* = 0.226, *S*.*E*. = 0.049, *p* < 0.001) while the dPMN did not (*β* = -0.042, *S*.*E*. =0.050, *p* = 0.402). When modeled separately, however, both the vPMN (*β* = 0.190, *S*.*E*. = 0.021, *p* < 0.001) and the dPMN (*β* = 0.170, *S*.*E*. = 0.022, *p* < 0.001) predicted Memory Quality (i.e., in models that included only one of the two paths). Fit statistics of alternate structural models (*Models 1-8*), which modeled a path from each region to Memory Quality after accounting for that region’s subnetwork participation, are presented in Table 3. Of the PM network regions, only the MPFC displayed a statistically significant region-specific ability to predict Memory Quality when controlling for its participation in its PM subnetwork (alpha = 0.05, FWE corrected for multiple comparisons). Inspection of the parameter estimates from this alternate model (*Model 4: MPFC*) suggests that the MPFC had a negative relationship with Memory Quality when controlling for its participation in the dPMN (*β* = -0.158, *S*.*E*. = 0.037, *p* < 0.001). The absence of other region-specific effects suggests that the contributions of the other PM regions were well described by the subnetwork level effects.

**Figure 2:**
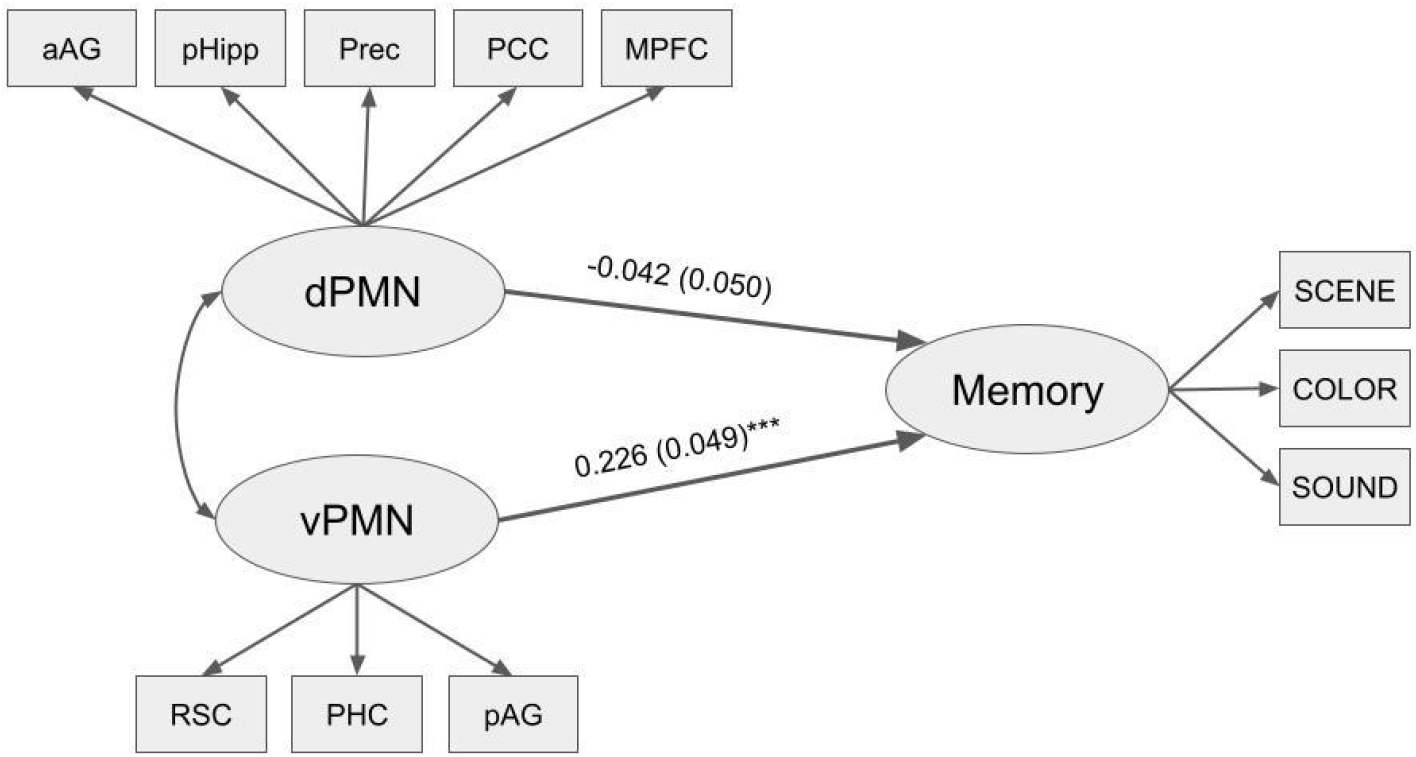
Structural Model. Path diagram representing the within-subject level of our two level baseline model (i.e., *Model 0*) with standardized parameter estimates (standard error in parentheses). See Figure 1 caption for abbreviations. * = *p* < .05, ** = *p* < .01, *** = *p* < .001.

**Table 4:** Structural Models. Table reports fit statistics for Models 0-8 as well as Wald’s Tests testing the constraint that the direct path from the ROI to the Memory Quality factor is zero. All Wald’s Tests p-values are Bonferroni corrected for multiple comparisons by multiplying each p value by 8 (the number of Wald’s tests performed). All p-values that were > 1 after this correction procedure were set to 1. Statistical tests report degrees of freedom in parentheses. Chi-square = chi-square test of exact fit, CFI = comparative fit index, TLI = Tucker–Lewis index, RMSEA = root mean square error of approximation, SRMR = standardized root mean squared residual.

| Title | Chi-square | Wald's Test | CFI | TLI | RMSEA | SRMR.Within | SRMR.Between |
| --- | --- | --- | --- | --- | --- | --- | --- |
| Model 0: Baseline | 534.781 (59), p: < 0.001 |  | 0.974 | 0.952 | 0.046 | 0.034 | 0.043 |
| Model 1: pHipp | 531.611 (58), p: < 0.001 | 3.86 (1), p: 0.395 | 0.974 | 0.951 | 0.046 | 0.034 | 0.043 |
| Model 2: Prec | 533.557 (58), p: < 0.001 | 6.176 (1), p: 0.104 | 0.974 | 0.951 | 0.046 | 0.034 | 0.043 |
| Model 3: PCC | 540.136 (58), p: < 0.001 | 0.575 (1), p: 1.000 | 0.974 | 0.951 | 0.046 | 0.034 | 0.044 |
| Model 4: MPFC | 525.826 (58), p: < 0.001 | 12.449 (1), p: 0.003 | 0.975 | 0.952 | 0.046 | 0.032 | 0.044 |
| Model 5: PHC | 563.181 (58), p: < 0.001 | 1.343 (1), p: 1.000 | 0.973 | 0.948 | 0.047 | 0.034 | 0.043 |
| Model 6: RSC | 533.996 (58), p: < 0.001 | 0.321 (1), p: 1.000 | 0.974 | 0.951 | 0.046 | 0.034 | 0.043 |
| Model 7: aAG | 531.518 (58), p: < 0.001 | 0.641 (1), p: 1.000 | 0.974 | 0.951 | 0.046 | 0.034 | 0.043 |
| Model 8: pAG | 545.002 (58), p: < 0.001 | 1.743 (1), p: 1.000 | 0.974 | 0.950 | 0.046 | 0.034 | 0.043 |

## Discussion

In the current study, we examined heterogeneity in the function of the PM network using a multilevel SEM framework. Our measurement models indicated that a two factor model with latent factors representing ventral and dorsal subnetworks was the best model for our neural data, compared to a single-factor model grouping all PM regions together. Our structural models indicated that the contributions of individual regions of the PM network to memory quality are largely subsumed by subnetwork-level contributions, with the exception of the MPFC which made a unique, region-specific contribution to memory quality. Interestingly, the region-specific contribution of the MPFC was found to be negative, such that less MPFC activation (when controlling for subnetwork membership) was associated with more accurate recollections.Together, these results reveal new insights into how memory outcomes can be explained by a combination of network-level and region-specific factors.

Our results support the presence of dissociable subnetworks within the PM network (Andrews-Hanna et al., 2010; Barnett et al., 2021; Buckner & DiNicola, 2019; Cooper et al., 2021). Previous studies have shown evidence for highly-related subnetworks during rest (Andrews-Hanna et al., 2010; Barnett et al., 2021) and during movie-watching (Cooper et al., 2021). Our results extend these findings, showing evidence that a similar subnetwork organization explains the trialwise involvement of PM regions during retrieval of multi-feature events. Our SEMs also showed that the coactivation of the vPMN makes contributions to memory quality that go above and beyond those made by coactivation of the dPMN (see Figure 2). The vPMN has previously been shown to modulate its connectivity in response to event transitions, and individual differences in episodic memory ability have been linked to dynamic changes in vPMN connectivity during movie watching (Cooper et al., 2021). The vPMN regions have also been shown to represent similar information during a memory-guided decision making task (Barnett et al., 2021). The vPMN is strongly related to episodic retrieval and autobiographical remembering, while portions of the dPMN have been linked to mentalizing about the mental states of others (Andrews-Hanna, Saxe, et al., 2014; Andrews-Hanna, Smallwood, et al., 2014). Additional evidence suggests that the vPMN may be particularly responsive to remembering and orienting towards space and the dPMN towards people (Peer et al., 2015; Silson et al., 2019). The fact that the vPMN in our dataset was uniquely related to memory quality could be reflective of our experimental design, which required the recollection of visual-spatial contextual details. In contrast, though the dPMN was strongly interconnected with the vPMN, it did not make an independent contribution to memory outcomes.

When taking into consideration the covariance among PM network regions, we did not find much evidence for independent, region-specific contributions, suggesting that the network-level effects could adequately account for their roles in predicting memory outcomes. Nevertheless, we had expected that there might be more region-specific effects, based on evidence that many of these regions play specialized roles in recollection. There are several reasons for why we did not see region-specific effects that we had hypothesized. One possibility is that there is something unique about our experimental paradigm that did not allow us to observe region-specific contributions. For example, the hippocampus may have emerged as making a region-specific contribution if we had operationalized our measure of memory success to more specifically target the hippocampus’ proposed function. In our analysis, we estimated a latent variable representing overall memory success from our feature specific memory measures and this latent measure served as the outcome variable. The hippocampus’ contribution to predicting this overall memory success measure may be subsumed by the network level contribution, but this may not be the case if the measure was more specific to successful pattern completion. Another possibility lies in how we modeled the neural response. In the current report, we modeled the neural response by assuming that it was transient, starting at the presentation of the memory cue during our ‘remember’ periods. Previous research suggests that the memory-related neural response in the angular gyrus is not transient with respect to the onset of recall, but is instead sustained throughout the duration of the recall period (Vilberg & Rugg, 2012, 2014). It is possible that modeling a sustained response throughout the recall period would allow for the identification of region-specific contributions of the angular gyrus. To test this possibility, all of our models were rerun using single trial estimates modeling the entire 4-second retrieval period. The key results of the current report remained unchanged in this model. Another possible explanation is that the identification of region-specific contributions within our framework assumes that the operations and representations of individual regions are not highly correlated. For example, it could be the case that the hippocampus and angular gyrus are, in reality, performing separable functions but those functions are so highly correlated in our task (e.g., because the functions of the angular gyrus are dependent on the output of the hippocampus) that they are well characterized by the network-level effect.

The one region in which we found a region-specific effect was the MPFC. The MPFC has been commonly described as part of the PM network (Ritchey & Cooper, 2020; Rugg & Vilberg, 2013) and is thought to support the formation and retrieval of schema-based representations (Schlichting & Preston, 2015; van Kesteren et al., 2012). In the current study, however, the MPFC was weakly correlated with the quality with which events were remembered, but was still positively correlated with other regions of the network (see Table 1). In recognition of its general participation in the PM network, we included the MPFC in our models. Our results indicated that, after controlling for MPFC’s participation in the dPMN subnetwork, the MPFC had a region-specific *negative* relationship with memory quality. However, we hesitate to draw strong conclusions regarding this finding. Previous reports suggest that MPFC activation is often *positively* correlated with measures of subjective memory success (H. Kim, 2016; McDermott et al., 2009; Spaniol et al., 2009). In the current study, because MPFC was only weakly correlated with memory quality, compared to the other regions in the analysis, the observed negative region-specific relationship might be explained by a statistical artifact. Specifically, the result seen here could be the result of a conditioning-on-a-collider bias, also known as *Berkson’s paradox* (Berkson, 1946; Lübke et al., 2020). In this paradox, two variables that, in reality, do not have a statistical association are induced to have a negative association by statistically controlling for a variable that they both cause. In the current scenario, it could be the case that MPFC activation and Memory Quality are (at least in part) correlated with increases in PM network coactivation, but Memory Quality and MPFC activation are not related to one another.The SEM methodology applied in the current report has a number of distinct advantages.Firstly, the current SEM approach has an advantage over previous reports of brain-behavior correlations in that it can simply and simultaneously capture the network-level and region-specific contributions of brain regions to behavioral phenomena. Second, the current report expanded upon previous deployments of this methodology (Bolt et al., 2018) by applying a multilevel SEM to simultaneously model within-subjects and between-subjects variation in the BOLD response, seeking to relate trial-by-trial, within-subjects variability in BOLD response to trial-by-trial variability in memory while controlling for individual differences. Thirdly, our dataset has a distinct advantage over previous studies of episodic remembering because it incorporates multiple measures of the quality of retrieval of an episode. This allowed us to model overall memory quality as a latent variable loading onto our measures of memory for 3 different features of each episode. By operationalizing memory success in this way, we were able to capture trial-to-trial variability in the joint remembering of event features. This is key, because holistic recollection is thought to be a key characteristic separating episodic remembering from other forms of memory (Tulving, 1983).

Our SEM approach is related to, but distinct from, other methods for relating regions and networks to episodic remembering. For example, previous studies have used data-driven, hierarchical clustering methods to parcellate PMN subnetworks (Andrews-Hanna et al., 2010; Barnett et al., 2021; Cooper et al., 2021), but did not relate trialwise coactivation within those subnetworks to episodic remembering. Another set of related methodological approaches is effective connectivity approaches. Specifically, some effective connectivity approaches also use SEM, but they use SEM to attempt to test hypothetical models of the underlying causal relations amongst regions of interest (e.g., McIntosh & Gonzalez-Lima, 1994). The latent variable modeling approach applied here, in contrast, does not attempt to make such causal inferences.Instead, our approach uses a latent variable to capture the coactivation seen within a network and relates this coactivation to a behavioral variable of interest. Lastly, the current approach is conceptually similar to partial least squares (PLS) analyses (Krishnan et al., 2011; McIntosh et al., 1996; McIntosh & Lobaugh, 2004). PLS involves maximizing the covariation between signal extracted from voxels of the brain and behavior, extracting latent variables reflecting distributed coactivation across the brain that explains variance in some behavior of interest. The SEM approach used in the current report is similar to PLS in that it also estimates a latent variable using the covariation of regional activation profiles, but has the advantage of being exclusively hypothesis driven and computationally and conceptually simpler. Many PLS applications (but not all Krishnan et al., 2011), in contrast, are data driven in nature. Additionally, PLS typically operates on all of the voxels collected during the course of an experiment, whereas the current approach operates on a set of hypothesized ROIs.

The current report makes an important contribution to the literature on the role of the PM network in episodic remembering. It does, however, have its limitations. Our multilevel approach allowed us to model trialwise neural actictivation and behavioral profiles while controlling for individual differences. Multilevel SEM, however, also allows researchers to build models of individual differences in neural activation and behavior beyond simply controlling for this important source of variability. We did not attempt to model individual differences in the current report in large part because our dataset would be underpowered to do so. Future research could utilize larger sample sizes to model individual differences related to particular participant characteristics (see Bolt et al., 2018 for an SEM application to individual differences). Additionally, the current analysis was focused on a set of *a-priori* ROIs that were the same across individuals. Although this is a good starting point and is a strategy often adopted by researchers, recent research in high-precision functional mapping suggests that individually defined ROIs may provide more accurate insights into network organization and function (Buckner & DiNicola, 2019; Gilmore et al., 2021). Finally, although our memory measures captured multiple aspects of each episode (specifically, memory for multiple episodic features), they may not have adequately captured the functioning of core alliances within the PM network (Ritchey & Cooper, 2020).

In conclusion, the brain is simultaneously composed of large scale brain networks and individual regions composing those networks. Here we demonstrate the importance of considering both network and regional levels of analysis when studying brain-behavior relationships, finding evidence in favor of a specific subnetwork organization of the PM network in its relation to episodic memory outcomes. Future work should continue to characterize the PM network by examining how these levels of analysis differentially support the various subprocesses and representations underlying episodic recollection.

## Acknowledgments

This work was supported by NIH grant R00MH103401 (M.R.) and NSF grant BCS-2047415 (M.R.). The data were collected at the Harvard Center for Brain Science, involving the use of instrumentation supported by the NIH Shared Instrumentation Grant Program; grant number S10OD020039.

## Footnotes

1 In multilevel models, SRMR at the between-cluster level may be large because the observed variances are small, not necessarily because there is a large extent of misfit.

